# An automated image analysis pipeline for wide-field optical redox imaging of patient-derived cancer organoids

**DOI:** 10.1101/2025.10.29.685427

**Authors:** Angela Hsu, Kayvan Samimi, Amani A. Gillette, Shirsa Udgata, Alexa E. Schmitz, Wenxuan Zhao, Dustin A. Deming, Melissa C. Skala

## Abstract

Wide-field optical redox imaging provides a fast and accessible method to monitor metabolic changes in cells and has recently been developed for drug screening in patient-derived cancer organoids (PDCOs). However, manual analysis of wide-field optical redox images is inefficient and laborious for large-scale drug screens. Here, we developed an automated pipeline for PDCO segmentation, single-PDCO tracking, and background correction in autofluorescence images. This pipeline was tested on two imaging systems over a 3-day time-course with two drug doses to demonstrate generalizability across imaging systems. Segmentation was performed using a fine-tuned Cellpose model, which when compared to manual masks, achieved mean Dice scores >0.8 across systems, indicating high reproducibility. Automated single-PDCO tracking was compared to manual tracking and the accuracy of the tracking algorithm exceeded 94% by two metrics, recall and Jaccard index. For background correction, the automated pipeline uses the full field-of-view to reduce sampling bias. Compared to the manual analysis pipeline, the automated pipeline resolves single-PDCO responses with comparable sensitivity to drug treatment but with over 127× faster processing time. This novel automated image analysis pipeline improves throughput and robustness in PDCO image analysis, which increases the accessibility and scalability of wide-field optical redox imaging for PDCO drug screening.

## Introduction

Colorectal cancer (CRC) is responsible for more than 50,000 deaths in the US annually^1^. CRC is a genetically and metabolic heterogeneous disease, but current treatments cannot effectively address patient and tumor heterogeneity^2^. There is differential sensitivity between patients. Within the same patient, tumors also respond to drugs differently. In addition to genetic heterogeneity, metabolic heterogeneity is apparent in CRC and contributes to drug resistance^3^. This emphasizes the need to develop a widely accessible and high-throughput drug screening method to understand the relationship between CRC heterogeneity and treatment response.

PDCOs are three-dimensional *in vitro* models derived from patient tumors that provide an excellent method to study subclonal resistance and metabolic heterogeneity^4–6^. PDCOs capture not only driver alterations but also subclones from the original tumor that drive secondary or acquired resistance^6^. PDCO metabolism provides an early measurement of treatment response compared to PDCO size, cell proliferation, and cell death^7–10^. However, existing PDCO metabolic assays have limitations in evaluating single PDCOs, longitudinal tracking in time course studies, and maintaining organoid integrity^11–20^. Therefore, a method that can provide repeated measurements of single PDCO metabolism without altering the PDCO culture could provide valuable insight into CRC treatments that address tumor heterogeneity.

Optical redox imaging addresses these limitations by monitoring single-PDCO metabolic heterogeneity over time without compromising the organoids with labeling or destructive techniques. Optical redox imaging leverages the autofluorescent properties of metabolic coenzymes nicotinamide adenine dinucleotide (phosphate) and flavin adenine dinucleotide (NAD(P)H and FAD) to measure the optical redox ratio (ORR), or the oxidation-reduction state of a cell. Here, NADH and NADPH are collectively referred to as NAD(P)H due to their overlapping spectral properties^21^. The ORR of PDCOs is often measured with multi-photon microscopy to resolve single cells, but multiphoton microscopy is costly, requires lengthy imaging times, and is therefore impractical for high-throughput drug screening^22^. Alternatively, wide-field microscopes are more affordable and accessible than multiphoton microscopes, and optical redox imaging with widefield microscopy only requires standard components such as a broadband excitation source, scientific monochrome camera, and standard 4′,6-diamidino-2-phenylindole (DAPI) and fluorescein isothiocyanate (FITC) filter cubes to collect NAD(P)H and FAD autofluorescence, respectively. Our group and others have used wide-field optical redox imaging at both cellular and tissue levels to evaluate oxidative stress, mitochondrial function, and drug responses^23–25^. Our prior work has demonstrated that wide-field optical redox imaging is a useful and nondestructive tool to assess patient treatment response by monitoring longitudinal metabolic changes in PDCOs at the single-PDCO level^26,27^. However, analysis of widefield optical redox imaging currently relies on a manual image analysis pipeline that is labor intensive, time consuming, and can be affected by human fatigue.

Here, we develop and optimize an automated image analysis pipeline for widefield optical redox imaging, which greatly accelerates drug screens of PDCOs while further improving the accessibility of this technique. To develop this automated pipeline, we first addressed challenges with automated PDCO segmentation including variability in PDCO shape, size, and focal plane positioning. Although prior studies have developed open source software to segment PDCOs, mostly based on bright field images, these solutions are not transferrable to autofluorescence images due to differences in contrast and signal to noise ratio (SNR)^28–31^. Second, PDCO variability requires tracking treatment response over time within the same PDCO to increase sensitivity to drug treatment compared to pooled PDCO analysis^26^. The automated PDCO tracking technique greatly improves the analysis time and reliability over manual PDCO tracking. Third, the current approach to calibrate each optical redox image requires a manual background region-of-interest (ROI) selection; whereas the automated background selection removes this additional manual step while improving the repeatability of the background measurement. Overall, we have developed automated modules for PDCO segmentation, tracking, and optical redox imaging background normalization to complete a fully automated image analysis pipeline for wide-field optical redox imaging that improves the throughput, accessibility, and reliability of PDCO drug response measurements. We demonstrate this pipeline across two separate imaging systems to highlight the generalizability of this approach, using a 3-day imaging time-course of PDCOs in response to two doses of drug treatment.

## Methods

### CRC PDCO isolation and culture

All work was conducted with Institutional Review Board (IRB) approval, and informed written consent was obtained from the patient through the University of Wisconsin (UW) Molecular Tumor Board Registry (UW IRB#2015-1370) or the UW Translational Science BioCore (UW IRB#2016-0934). All methods were performed in accordance with the relevant guidelines and regulations. CRC tissue was obtained via primary resection and processed as previously described^32,33^. Processed PDCO suspension was immediately mixed in a 1:1 ratio with Matrigel (Corning) and conditioned media. Droplet suspensions were plated, set for 3-5 minutes at 37°C, and then inverted for at least 30 minutes to solidify the matrix and avoid direct contact of PDCOs with the interface. Plated cultures were overlaid with 450 µL of media, consisting of DMEM/F12 (Invitrogen) supplemented with 1x Glutamax (Invitrogen), 10 mM HEPES (Fisher), 50 IU/mL penicillin-streptomycin (Invitrogen), 50 ng/mL EGF (Invitrogen), and mixed 1:1 with WNT3a-conditioned media^33^. PDCOs were incubated at 37°C in 5% CO₂ and media was replaced every 48–72 hours. A single CRC PDCO line was used for this study.

### Romidepsin treatment experiment

PDCOs were collected from 24-well culture plates and transferred to 24-well glass bottom plates (Corning), then rested for 24 hours. Pre-treatment (day 0) images of the PDCOs were acquired using a Nikon Ti-S inverted microscope or Keyence BZ-X810 microscope as described below. Romidepsin (MedChem Express; HY-15149) was prepared at 10 mM in DMSO and diluted to 30 or 100 nM in culture media, while no DMSO was added to the control media. PDCOs were treated with control media, 30 nM romidepsin, or 100 nM romidepsin for 48 hours, and imaged again at 24 hours (day 1) and 48 hours (day 2).

### Wide-field optical redox imaging

This study performs wide-field optical redox imaging with two imaging systems (Nikon and Keyence) to demonstrate the generalizability of our automated image analysis pipeline.

#### Nikon

A Nikon Ti-S microscope coupled with a SOLA light engine (300 to 650 nm, Lumencor, Beaverton, Oregon, United States) was used. Images were acquired using NIS Elements software (Nikon), a 4x air, 0.13 NA objective (Plan Apo, Nikon), and a Hamamatsu Flash4 digital CMOS camera (Hamamatsu City, Japan). NAD(P)H was excited for 3 seconds through a 360/40 nm filter at 40% power (0.25 mJ/cm²), and emission was collected using a 400 nm dichroic mirror and a standard 4′,6-diamidino-2-phenylindole (DAPI, 460/50 nm) filter. FAD was sequentially excited for 3.5 seconds through a 480/30 nm filter at 40% power (0.531 mJ/cm²), with emission collected using a 505 nm dichroic mirror and a standard fluorescein isothiocyanate (FITC, 535/20 nm) filter. Four fields of view (FOV) (2048 x 2048 pixels; 1.61 μm/pixel) of each PDCO preparation treated with control media, 30 nM romidepsin, or 100 nM romidepsin were imaged pre-treatment (day 0) and post-treatment at 24 hours (day 1) and 48 hours (day 2).

#### Keyence

A Keyence BZ-X810 microscope (Osaka, Japan) coupled with a 40 W LED (Osaka, Japan) using a 4x air, 0.20 NA objective was used. Images were acquired with a 2/3 inch, 2.83 million pixel monochrome CCD (Osaka, Japan). NAD(P)H was excited for 3 seconds at 20% light power through a DAPI filter cube (360/40ex, 400nm dichroic, 460/50em; Keyence BZ-X DAPI filter). FAD was sequentially excited for 3.5 seconds at 40% light power through a custom filter (480/40ex, 510 dichroic, 535/50em; Keyence BZ-X FITC filter). Each PDCO population treated with control media, 30 nM romidepsin, or 100 nM romidepsin was imaged pre-treatment (day 0) and post-treatment at 24 hours (day 1) and 48 hours (day 2). The same wells of PDCOs were imaged with both the Keyence and Nikon systems, however, on the Keyence system automatic stitching was performed with a 30% overlap between adjacent FOV, yielding one large FOV (3594 x 3486 pixels; 2.4 μm/pixel) per well.

### Manual Image Analysis Pipeline

#### Manual segmentation and leading-edge analysis

PDCO masks were manually segmented from NAD(P)H intensity images using a CellProfiler pipeline^27^. PDCOs consist of (from innermost to outermost) necrotic cores, quiescent zones, and proliferating zones. Proliferating zones at the outer edge of PDCOs are the most metabolically active and most sensitive to treatment^34,35^. To isolate redox changes in the proliferating zone, our group developed “leading-edge analysis” in a recent study^36^. Briefly, we calculate the radius of each single-PDCO mask, and subtract 20 pixels in Nikon or 13 pixels in Keyence (due to differences in resolution), roughly 32 µm, from the radius to generate a core mask, which is then subtracted from the single-PDCO mask to create a leading-edge mask. The normalized ORR and redox ratio changes were then calculated from leading-edge masks.

#### Sampled Background Normalization of ORR

To address issues such as variations in different imaging systems, instrumental drift within an imaging system, uneven illumination, and autofluorescence from Matrigel, ORR values were normalized to a background value within each image. In the manual pipeline, background value refers to the average fluorescence intensity of five PDCO-free regions selected by an individual (**Fig. 1**).

**Figure 1:**
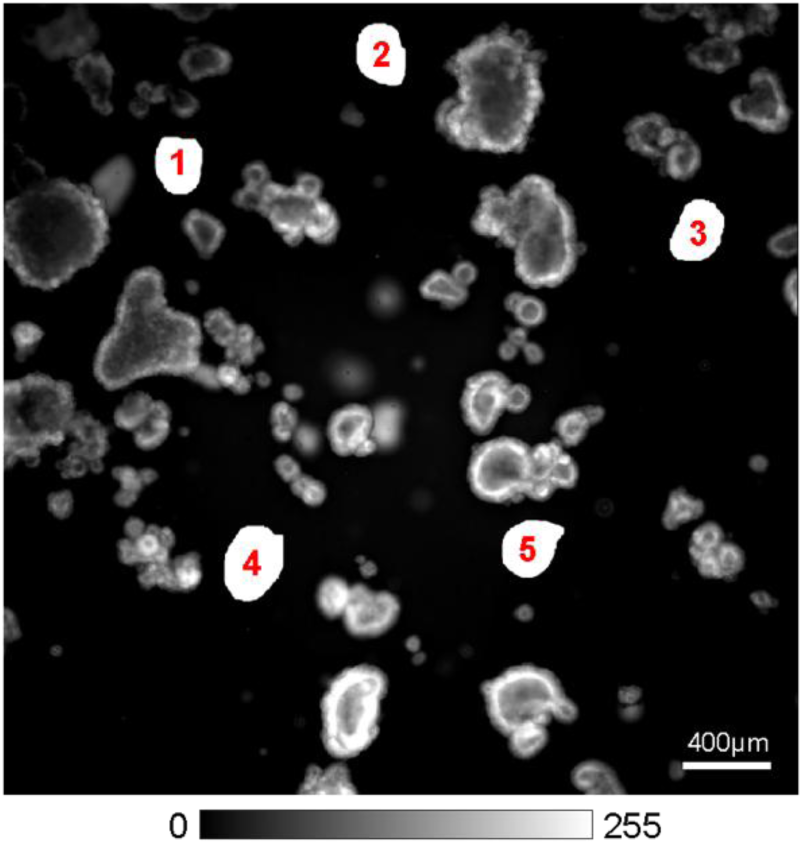
Representative NAD(P)H intensity image of sampled background normalization from the manual pipeline. An individual selects five PDCO-free regions (white) and uses the average fluorescence intensity as the background value to normalize NAD(P)H and FAD intensity images. Color scale bar reflects intensity scaled 0-255 for visualization. Numbered PDCO-free regions are illustrated in white for visualization only and have intensity values near zero.

The background value is then used to calculate the normalized ORR for each PDCO, defined as in Eq. 1-3:

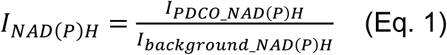

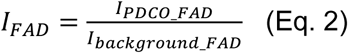

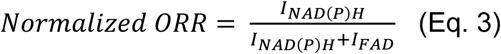

Where I_PDCO_NAD(P)H_ is the mean intensity of NAD(P)H across the leading-edge mask of the PDCO, and I_background_NAD(P)H_ is the mean intensity of the five PDCO-free regions across the entire NAD(P)H image. I_PDCO_FAD_ is the mean intensity of FAD across the leading-edge mask of the PDCO, and I_background_FAD_ is the mean intensity of the five PDCO-free regions across the entire FAD image.

For the manual pipeline, ΔORR is defined as the normalized ORR of an individual PDCO (Eq. 3) minus the average of all PDCOs in that condition measured on day 0 as in Eq 4:

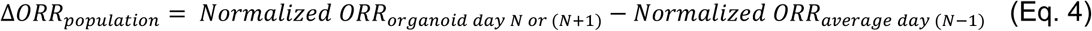

For generalizability, we denote consecutive days as day (N-1), day N, and day (N+1). In this study, they correspond to day 0, day 1, and day 2, respectively. Where “Normalized ORR_organoid day N_” is the background normalized ORR of individual PDCO on day 1, and “Normalized ORR_organoid day (N+1)_” is the background normalized ORR of individual PDCO on day 2. “Normalized ORR_average day (N-1)_” is the background normalized ORR of the same PDCO population measured on day 0 prior to treatment.

### Automated Image Analysis Pipeline

All scripts in the automated image analysis pipeline were written in Python 3.10.12.

### Algorithm overview: PDCO segmentation, longitudinal single PDCO tracking, and background correction

Prior work has demonstrated that wide-field optical redox imaging is a rapid, sensitive, and non-invasive method to measure treatment response and heterogeneity in PDCOs^26,36,37^. To meet the demands of analyzing large-scale drug screens, a more accessible and high-throughput image analysis method is needed. This study aims to address this need by automating PDCO segmentation, longitudinal single PDCO tracking, and background correction. A complete workflow of the improved analysis pipeline is demonstrated here **(Fig. 2A-F)**.

**Figure 2:**
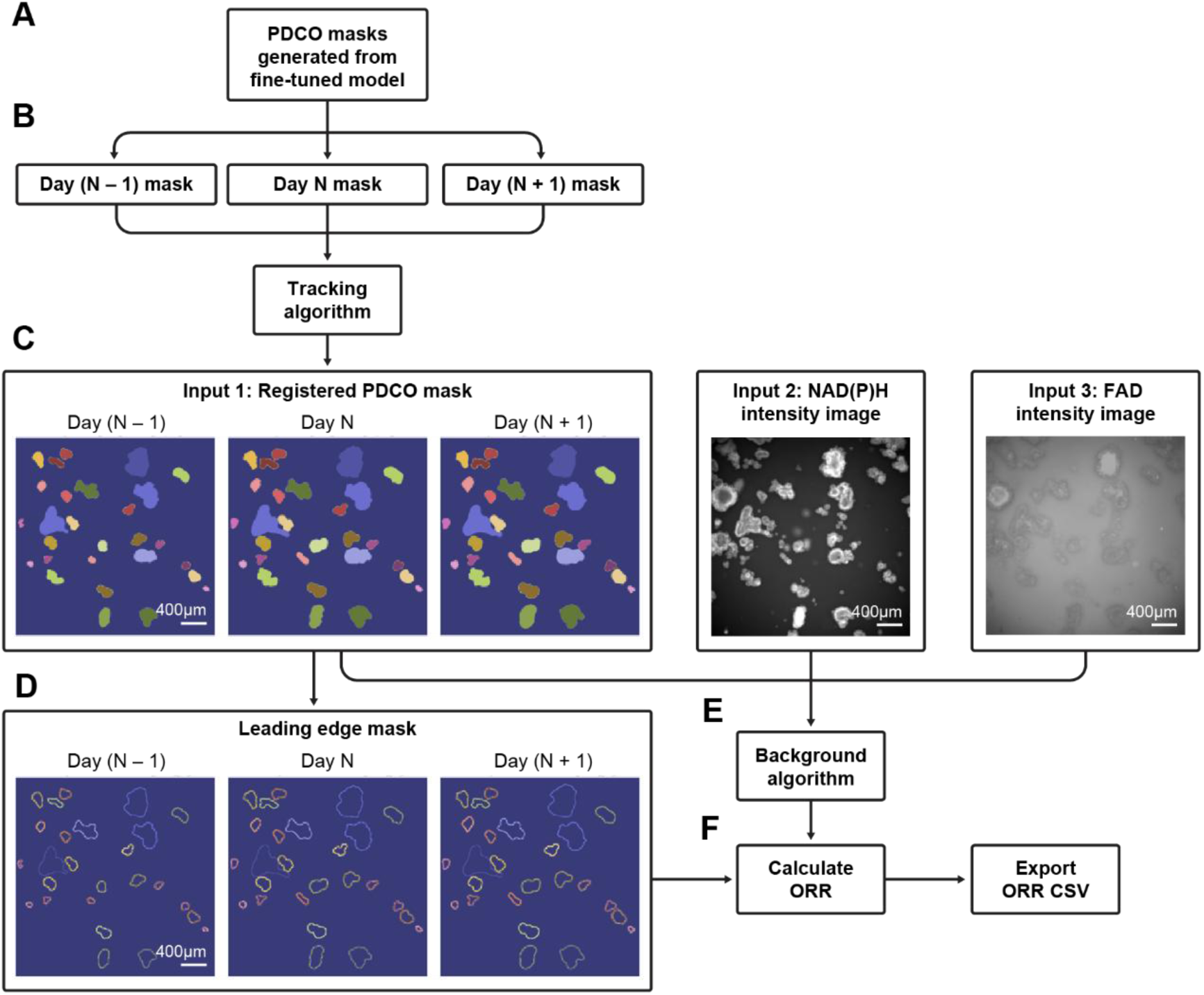
Workflow of the automated wide-field PDCO ORR image analysis pipeline. (A) The fine-tuned Cellpose model was used to generate new masks with a batch-processing script. A separate script is used to remove incomplete PDCO masks with pixels touching the borders of the ROI (not shown). (B) Day (N-1), day N, and day (N+1) masks can be used as inputs for the tracking algorithm, which assigns matching objects in each mask with the same unique ID based on cross-correlation scores and location. (C) Masks with matched unique IDs, with NAD(P)H and FAD intensity images, can then be used as inputs for the background pixel identification algorithm. (D) Leading-edge masks are created. (E) NAD(P)H and FAD intensity images are background normalized and used to generate the ORR image. (F) The ORR mean values are calculated in the leading-edge mask for each PDCO across timepoints and are saved to the output CSV file.

#### Automated Segmentation

Following the instructions described in "Cellpose 2.0: How to Train Your Own Model," the cyto3 model was fine-tuned using the following training parameters: pretrained_model cyto3, n_epochs 100, learning_rate 0.1, and weight_decay 0.0001^38,39^. 55 PDCO NAD(P)H intensity images and their manual masks were used to train the model. 15 NAD(P)H intensity images and their manual masks were tested, including 3 images of 30nM romidepsin treatment taken with the Nikon microscope. All images in the training and testing sets were taken with the Nikon microscope. Only whole PDCOs were masked in all of the images used for training and testing.

The fine-tuned Cellpose model was used to generate PDCO masks with a batch processing script. PDCOs with pixels touching the borders of the masks were removed by a post-processing script. Only whole PDCO masks were used for the tracking algorithm (**Fig. 3**) and background mask identification (**Fig. 4**) described below.

**Figure 3:**
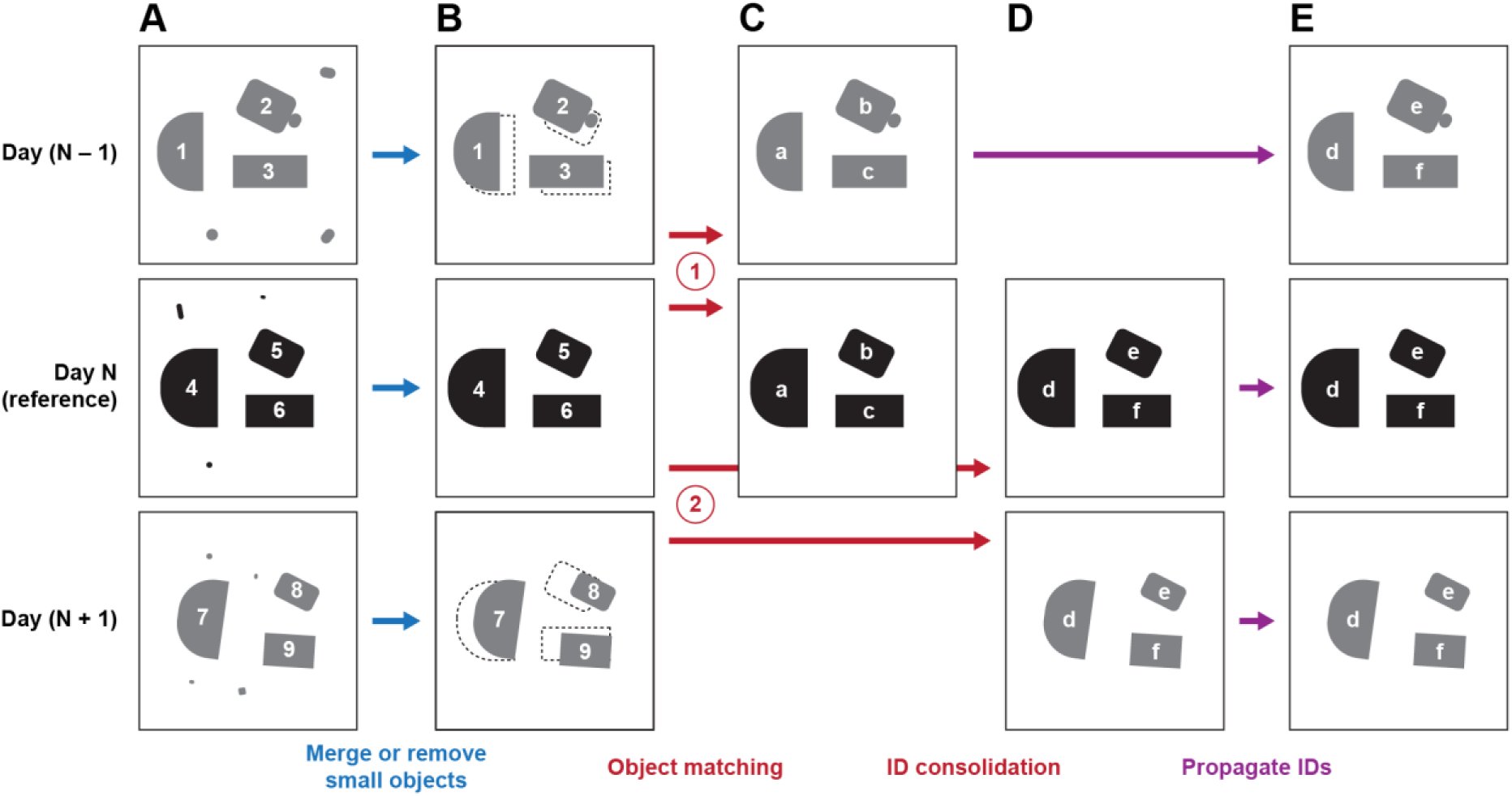
General workflow for the algorithm to track PDCOs within NAD(P)H images over multiple time-points. Coarse image registration occurs before the following steps (not shown). (A-B) Light blue arrows indicate the removal of small objects likely to be debris. (B-D) Red arrows indicate the object matching and ID consolidation step where the old IDs (numbers) are updated to new IDs (letters). (B-C) Red circled numbers indicate object matching based on cross-correlation, first between day (N-1) and day N, then day (N+1) and day N. (C-D) ID consolidation occurs between the two updated day N masks, linking ID pairs (*e.g.*, a d, b e, c f). (C-E) Purple arrows indicate paired IDs are propagated to ensure updated IDs are consistent across all days for each PDCO. Numbered and lettered shapes represent PDCOs. Numbers and letters represent IDs, or unique identification labels. Numbered IDs are unique to each PDCO in each image. Letter IDs are unique to each tracked PDCO across two or more time points. Changes in shape size and orientation from day (N-1) to day (N+1) illustrate changes in PDCOs after treatment.

**Figure 4:**
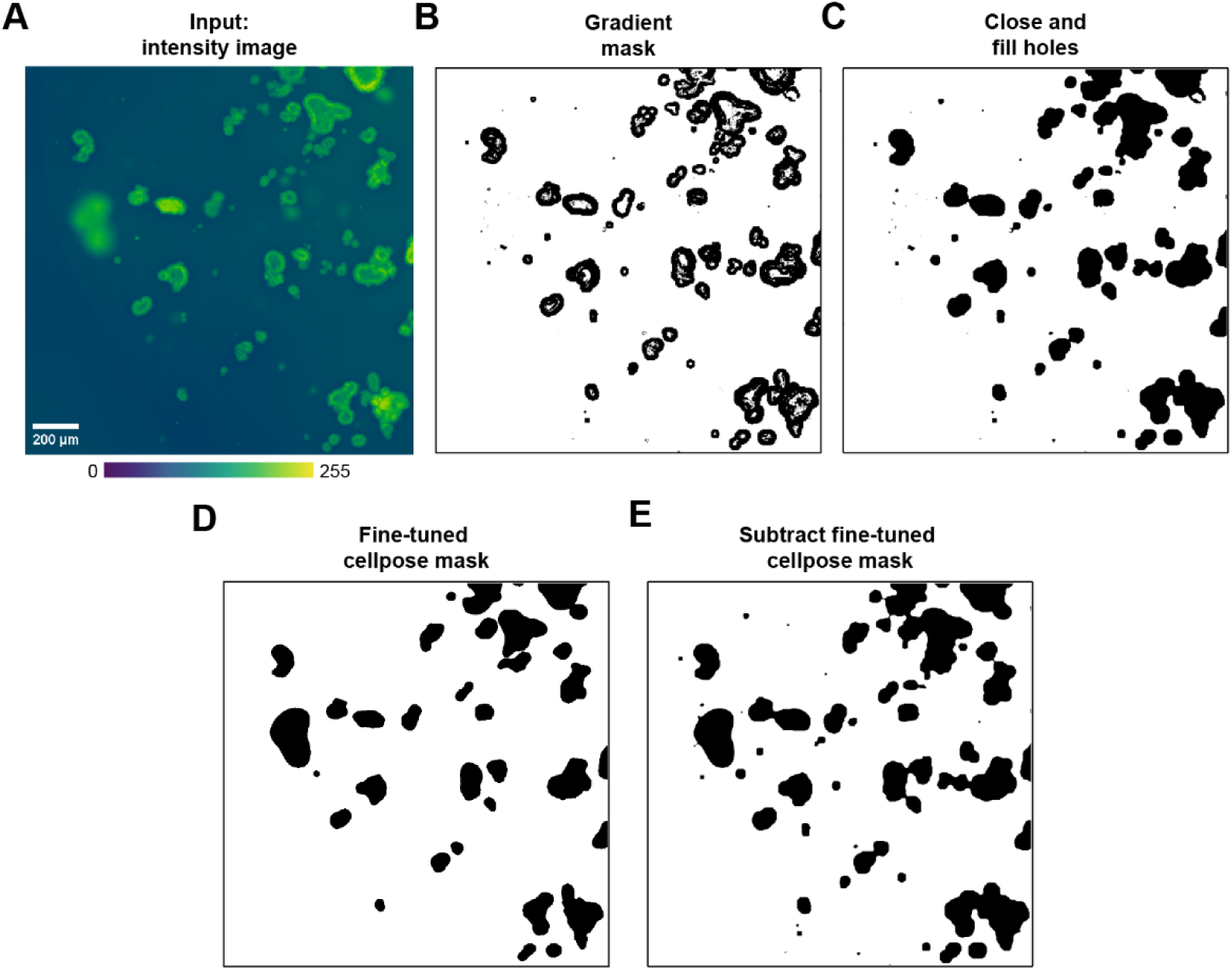
General workflow for the algorithm to automatically detect background pixels within autofluorescence images. (A) Representative input NAD(P)H intensity image. NAD(P)H and FAD images are separately processed to identify background pixels for each image. Color scale bar scaled to 0-255. (B) Gradient mask generated after Sobel operation. (C) Mask after applying closing-and-filling-holes operation. (D) Fine-tuned Cellpose mask (E) Final background mask generated after subtracting the area covered by the fine-tuned Cellpose mask.

#### Tracking Algorithm

The tracking algorithm **(Fig. 3)** used the following modules and libraries: glob, matplotlib, tifffile, opencv-python, SciPy, and scikit-image^40–42^. Here, a mask refers to a binary image of segmented PDCOs, and an object refers to an individual PDCO in the mask. Up to 3 masks (i.e., time points) can be input into the tracking algorithm but the program may be modified to accommodate more masks. Here, the workflow up until the ID consolidation step is illustrated with only the day (N-1) and day N image pair. The same workflow is repeated for the day N and day (N+1) image pair **(Fig. 3A-C)**. After ID consolidation, IDs obtained from the day N and day (N+1) pair are propagated to day (N-1) **(Fig. 3D-E)**.

First, the tracking algorithm preprocesses the masks by removing small objects below 1000 pixels **(Fig. 3A-B)**. Next, within an image time-series, day (N-1) mask is coarsely registered to day N mask. 2D cross-correlation scores are computed within an offset search area of ±25% of the mask size in X and Y directions. The translation vector resulting in the highest cross-correlation score between the two masks is identified. This vector is applied to coarsely register day (N-1) mask to day N mask within an image time-series.

To match each PDCO over time, objects in the day (N-1) mask are sorted by size to prioritize larger objects as they tend to have more prominent features. Each object in the day (N-1) mask is extracted as a template, and a search region (25% of the mask size) is determined in the day N mask. Each object in the search region is tested for a match with the template. A match is confirmed if the cross-correlation score exceeds a threshold dynamically scaled (between 0.2 and 0.5) based on the object’s size **(Fig. 3B-C)**. The scale of cross-correlation score here is normalized to 0-1. The threshold is defined as the low minimum acceptable cross-correlation score (0.2) plus the difference between the high (0.5) and low (0.2) minimum cross-correlation scores, scaled by 10 times object’s size relative to the image. This ensures small objects aren’t penalized for having lower correlation scores and requires larger objects with more prominent features to have higher correlation scores to match. Matched objects are then set to zero in the day N mask to prevent duplicate matching before the process is repeated to find matches for the remaining objects.

To address merging objects, labels of all objects in the mask are extracted and a perimeter mask is created by outlining each object’s contours. Neighbors with the most overlapping pixels are identified, and merging is performed if the sizes of both objects fall below 1% of the FOV and their overlapping perimeter exceeds 40%. The smaller object is merged into the larger object.

Once matching is complete, the masks are updated with new identification numbers (IDs) **(Fig. 3B-C)**. Each pair of matched objects is assigned the same ID. Consolidation of IDs is performed between the updated day N mask from matching with day (N-1) mask and the updated day N mask from matching with day (N+1) mask **(Fig. 3C-D)**. This step links ID pairs that identify the same PDCO in the reference day N mask.

Finally, the IDs from the day N-and-day (N+1) matching are propagated to replace old IDs in the day (N-1) mask, according to the ID pairings in the consolidation step **(Fig. 3D-E)**. Masks are updated with new IDs only for matched objects. Unmatched objects are removed from the masks.

With consistent IDs across day (N-1) and day (N+1) masks, ΔORR for each PDCO can then be calculated by subtracting the normalized ORR of an individual PDCO on day N-1 from the normalized ORR of the same PDCO on day N or (N+1) as in Eq. 5:

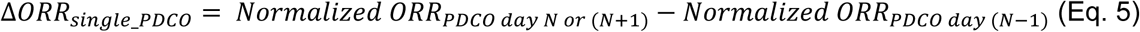

Where “Normalized ORR_PDCO day N or (N+1)_” is the background normalized ORR of individual PDCO on day N or (N+1), and “Normalized ORR_PDCO day (N-1)_” is the background normalized ORR of the same PDCO measured on day (N-1) prior to treatment, as in Eq. 3.

#### Image Background Mask Identification

It is inevitable that some PDCOs are not captured with the fine-tuned Cellpose masks. If a simple inverse mask was used, those PDCOs would be included in the background, skewing the accuracy of the background values used for normalization. Therefore, a background algorithm is developed.

The algorithm used the following modules and libraries: matplotlib, pandas, tqdm, natsorted, numpy, tifffile, opencv-python, SciPy, scikit-image^40–44^. The algorithm requires a fine-tuned Cellpose mask, NAD(P)H intensity image, and FAD intensity image to generate a background mask.

First, a gradient mask is generated from the intensity images **(Fig. 4A)** by calculating the normalized gradient magnitude using the Sobel operator and applying Otsu’s threshold to separate areas with high spatial intensity gradient, like PDCO edges, from the low-gradient background **(Fig. 4B)**. Next, the image is subsampled, keeping every fourth pixel in both the horizontal and vertical directions and these steps are repeated. This is done to increase the intensity gradients and allow detection of more homogenous PDCOs that might have been missed in the first round. While this operation highlights the PDCO edges, the PDCO interiors often have uniform areas of low intensity gradient that are not selected by this operation. Therefore, morphological closing and filling of holes is performed to capture the interior of PDCOs **(Fig. 4C)**. During the development of the algorithm, overmasking occurred frequently at this step. To detect potential overmasking, if the filled gradient mask exceeds 80% of the total image or if the largest object is five times larger than the second largest object, the gradient mask is reverted to the previous mask, before hole filling. The last background mask is then inverted and overlaid with the fine-tuned Cellpose mask **(Fig. 4D)** to produce a foreground mask **(Fig. 4E)** to ensure PDCOs of interest are correctly identified as foreground and excluded from the background mask.

Next, the top 2% brightest pixels in the image are added to the foreground mask and objects smaller than 25 pixels are removed from the foreground mask, and hole closing and filling operations are performed again. The final background mask is produced by inverting this updated foreground mask.

Finally, NAD(P)H and FAD background masks are combined to create the final background mask. If overmasking is detected when both masks are combined, the mask with fewer background pixels is selected. If no overmasking is detected, the NAD(P)H and FAD masks are combined using a logical AND operation.

In the automated pipeline, the average fluorescence intensity of the background mask pixels in the NAD(P)H image gives the background NAD(P)H value (I_background_NAD(P)H_) in Eq. 1, and the average fluorescence intensity of the background mask pixels in the FAD image give the background FAD value (I_background_FAD_) in Eq. 2, used to calculate the normalized ORR in Eq. 3.

#### Leading-edge masks

Leading-edge masks are only used to calculate final ORR (Eq. 3) and ΔORR_single PDCO (Eq. 5) values after tracking and background normalization. In the automated analysis pipeline, a leading-edge mask is created by first eroding a single-PDCO mask by 32 µm, or 20 pixels for Nikon images and 13 pixels for Keyence images due to differences in resolution, followed by subtracting the eroded mask from the original single-PDCO mask. This procedure is the same as the manual pipeline.

### Statistical Analysis

#### Dice Similarity Coefficient

The Dice Similarity Coefficient, or Dice Score, was used to calculate the similarity between the fine-tuned Cellpose masks and manual masks. The Dice score is defined as in Eq. 6:

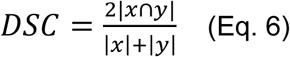

Where |*x*| is the number of pixels in the first mask, and |*y*| is the number of pixels in the second mask. |*x* ∩ *y*|is the number of pixels shared by both masks. The Dice score ranges from 0 to 1, where 0 indicates no overlap between two masks, and 1 indicates complete overlap.

#### Recall

Recall was used to assess the accuracy of the tracking algorithm. Recall is defined as in Eq. 7:

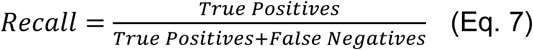

Where true positives are the shared tracks between manual tracking and automated tracking and false negatives are tracks unique to manual tracking. The recall metric measures how many of the manual tracks are correctly detected by the tracking algorithm.

#### Jaccard index

Jaccard index was also used to assess the accuracy of the tracking algorithm. Jaccard index is defined as in Eq. 8:

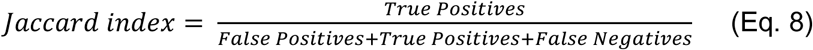

Where true positives are the shared tracks between manual tracking and automated tracking, false positives are tracks unique to automated tracking, and false negatives are tracks unique to manual tracking. The Jaccard index metric measures how many of the unique tracks found in either manual or automated tracking are correctly detected by the tracking algorithm.

#### Median Glass’s Delta (mG*Δ*)

mGΔ was used to measure the effect size of romidepsin treatment because it provides a more conservative assessment of treatment effects in large sample sizes compared to ANOVA.

mGΔ is defined as in Eq. 9 and 10:

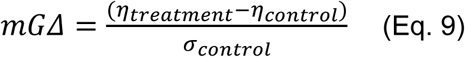

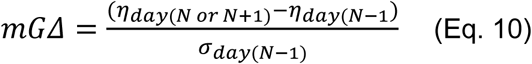

Where η_treatment_ is the median ΔORR (Eq. 4 or 5) of the treatment group, η_control_ is the median ΔORR of the control group, and σ_control_ is the standard deviation of the ΔORR of the control group. In manual analysis, ΔORR_population_ (Eq. 4) was used. In automated analysis, ΔORR_single_PDCO_ (Eq. 5) was used.

For Eq. 10, where η_day (N-1)_ is the median of ORR (Eq. 3) of the day 0 group, η_day N_ is the median of ORR (Eq. 3) of the day 1 group, and η_day (N+1)_ is the median of ORR (Eq. 3) of the day 2 group. σ_day (N-1)_ is the standard deviation of the ORR of the day 0 group.

#### Percent Difference

Percent difference in ORR between the manual and automated methods was calculated. Percent difference is defined as in Eq. 11:

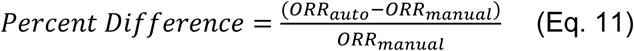

Where ORR_auto_ is the ORR obtained from the automated pipeline, and ORR_manual_ is the same ORR obtained from the manual pipeline.

## Results

### Similarity of fine-tuned Cellpose masks and manual masks

In prior work, PDCOs were segmented manually using NAD(P)H intensity images in CellProfiler. Manual segmentation is time-consuming and prone to human error, especially when analyzing images on a large scale. Therefore, we adopted Cellpose, a deep-learning based, generalist algorithm, as an automated segmentation tool^38^. However, existing Cellpose models performed poorly with PDCO segmentation due to their complex, heterogeneous morphology that lacks consistent shapes and features for easy identification^36, 37^. In addition, wide-field autofluorescence images have low SNR, making segmentation challenging^45^. To address these issues, we fine-tuned the cyto3 model using 55 NAD(P)H intensity images and their corresponding masks created manually^39^. Fifteen NAD(P)H intensity images and their manual masks were tested, including 3 images of 30 nM romidepsin treatment taken with the Nikon microscope. A batch processing script using this fine-tuned model then generates PDCO masks, and PDCOs at image borders are removed afterwards. Treatment with romidepsin was chosen for this study to evaluate optical redox imaging’s ability to assess new CRC therapies. Romidepsin is a histone deacetylase (HDAC) inhibitor FDA-approved for cutaneous T-cell lymphoma, peripheral T-cell lymphoma, and multiple myeloma.^46–48^ It has previously been shown to inhibit tumor growth in CRC tumor models^49,50^

Manual and fine-tuned Cellpose masks demonstrated strong agreement when overlaid for comparison **(Fig. 5A, 5B)**. In the few instances automated and manual masks disagree, PDCOs are out of focus or located at the image borders. The fine-tuned model often includes out-of-focus PDCOs as part of the mask, whereas a trained individual excludes them. Interestingly, despite the training dataset excluding PDCOs at the borders, the fine-tuned model consistently incorporates all PDCOs at the edges of the image. We therefore removed these PDCOs at the image edges using a post-processing script.

**Figure 5.**
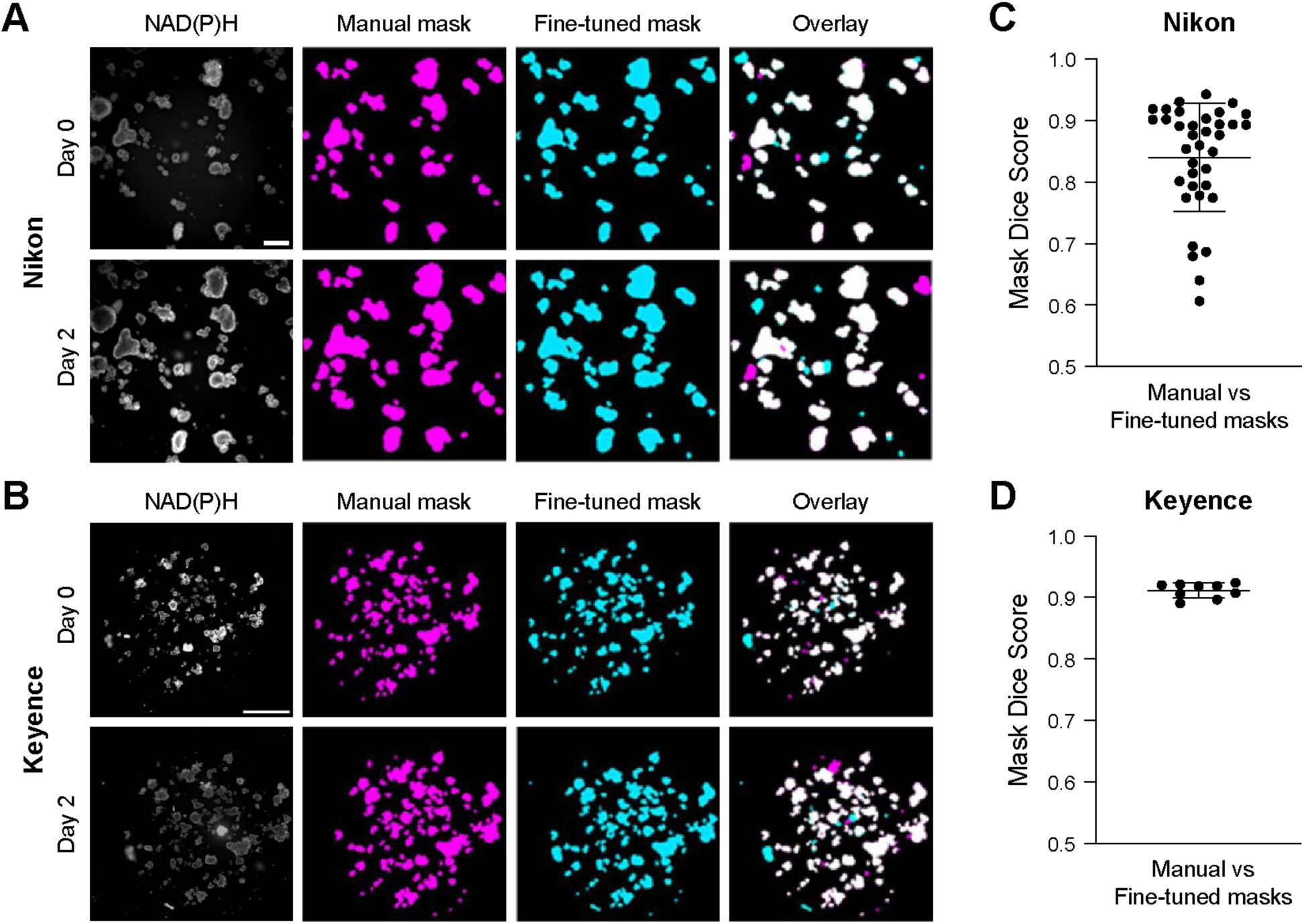
Dice score measurements reveal high agreement between manually segmented masks and Cellpose fine-tuned masks. Representative images of wide-field NAD(P)H intensity images of CRC PDCOs before (day 0) and 2 days after 30 nM romidepsin treatment (day 2) for (A) Nikon microscope and (B) Keyence microscope images. Masks were overlaid for manually segmented (magenta) and Cellpose fine-tuned masks (cyan) where more overlap (white) results in a higher dice score. The Dice Similarity Coefficient, or Dice Score, is computed by taking twice the white pixel area and dividing it by the combined pixel areas of the magenta and cyan masks (Eq. 6). Nikon scale bar = 400 µm. Keyence scale bar = 2000 µm. The same wells of PDCOs were imaged with both the Keyence and Nikon systems, however, on the Keyence system automatic stitching was performed with a 30% overlap between adjacent FOV, yielding one large FOV per well. For (C) Nikon Dice Score, each dot represents a FOV, 4 acquired per PDCO well (*n* = 36). For (D) Keyence Dice Score, each dot represents a full-field image of the same PDCO well (*n* = 9). Center line is mean. Error bars are standard deviation.

To quantitatively validate the fine-tuned Cellpose masks, a Dice Similarity Coefficient, or dice score, is used to measure the similarity between the manual and automated masks **(Fig. 5C, 5D)**. To demonstrate the generalizability of our approach, images of the same PDCO dishes were acquired on the same day with both Nikon and Keyence microscopes, which have different field-of-view dimensions and image resolutions. The Keyence microscope performs automatic stitching of fields of views and therefore captures a larger total area than the Nikon microscope. Both imaging systems achieved mean dice scores exceeding 0.8. The mean dice score for the Nikon microscope (0.84) is somewhat lower than that of the Keyence microscope (0.91), perhaps due to fewer PDCOs in the Nikon images (corresponding to its smaller area images). Overall, this indicates the high reproducibility of our fine-tuned model across different imaging systems, making this a suitable segmentation method for our high-throughput and accessible analysis pipeline.

### Tracking algorithm captures single-PDCO redox changes over time

Single-PDCO analysis provides an opportunity to study the impact of tumor heterogeneity on drug resistance. Prior studies have shown that single-PDCO tracking also shows higher sensitivity to treatment response compared to pooled analysis, emphasizing the need for reliable methods to track single PDCOs in a high-throughput drug screen^4,26^. To address this need, the second innovation in this automated analysis pipeline is an algorithm designed to track single PDCO redox changes across different time points. Two metrics (recall and Jaccard index, Eq. 7 and 8, respectively) were used to test the accuracy of the tracking algorithm, and both metrics show >94% accuracy rate **(Fig. 6A-B)**.

**Figure 6:**
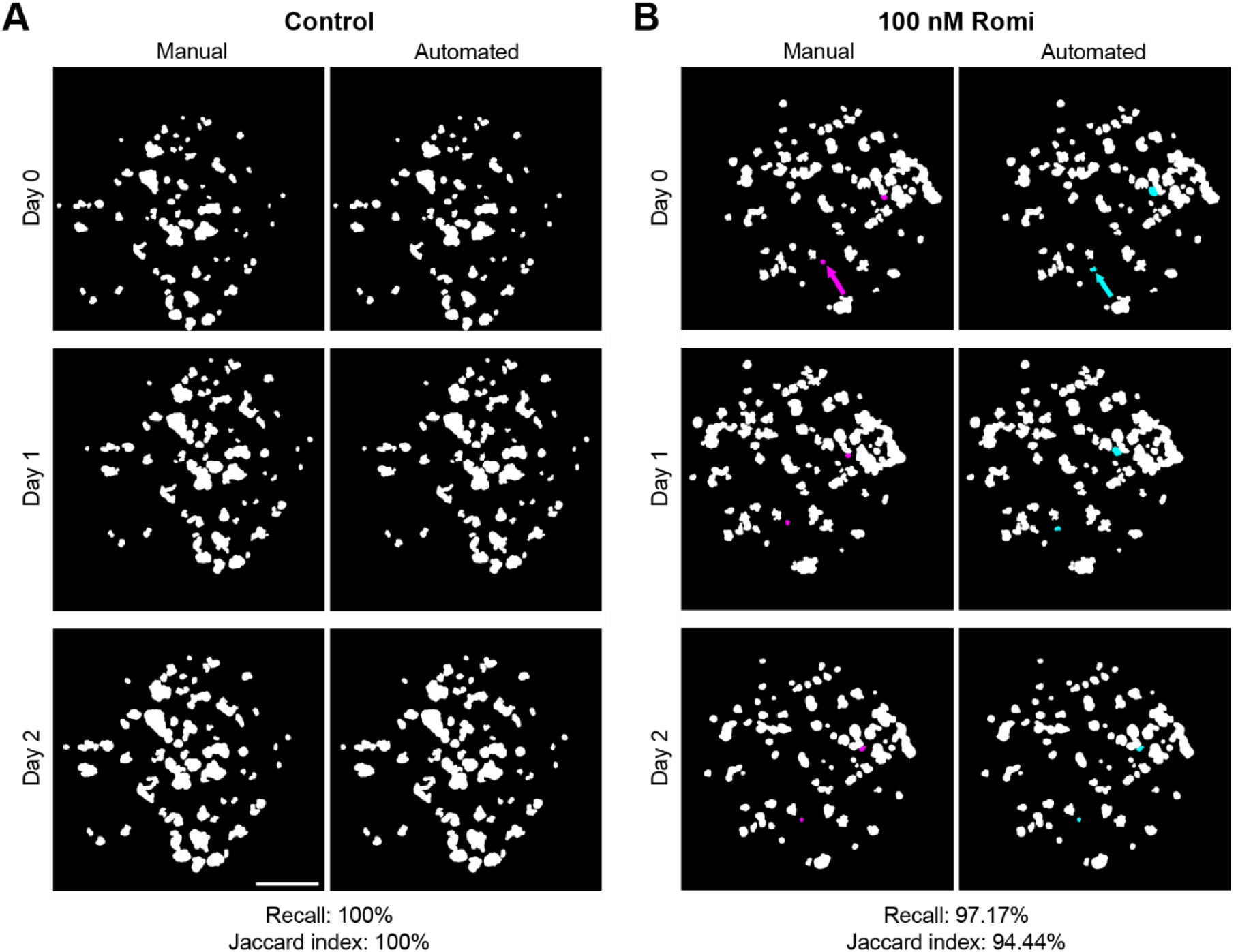
Both recall and Jaccard index show high accuracy for the tracking algorithm compared to manual tracking. 3-day imaging time course of CRC PDCOs treated with (A) control media or (B) 100nM romidepsin acquired with the Keyence microscope. White represents tracks shared by both manual tracking and automated tracking. Magenta represents tracks unique to manual tracking. Cyan represents tracks unique to automated tracking. For Control: Shared tracks n = 69. Unique manual tracks n = 0. Unique automated tracks n = 0. For 100 nM romidepsin: Shared tracks n = 68. Unique manual tracks n = 2. Unique automated tracks n = 2. Scale bar = 2000 µm.

Color-coded representative Nikon images illustrate how unique IDs are assigned to each PDCO consistently in images acquired on different days **(Fig. 7A)**. To demonstrate and validate single-PDCO tracking with redox values, we first manually segmented, tracked, and calculated the normalized ORR (Eq. 3) of each PDCO on days 0, 1, and 2 of romidepsin or control treatment **(Fig. 7B)**. Only Keyence images were used in this part of the study. For manual tracking, unique IDs were assigned to PDCOs in each image according to the order in which they were segmented, which means that IDs of the same PDCO are not consistent across images acquired on different days. Then, a manual process of matching redox ratio values of PDCOs across days was performed. Next, we automatically segmented, tracked, and calculated the normalized ORR (Eq. 3, **Fig. 7B)**. We then calculated the percent difference between the ORR obtained from the manual pipeline and the same ORR obtained from the automated pipeline for each individual PDCO (Eq. 11). The percent difference in ORR between the two methods is <25% for all treatment groups **(Fig. 7C)**. mGΔ for all groups are also similar between two pipelines. This suggests there is high agreement between the automated and manual pipelines.

**Figure 7.**
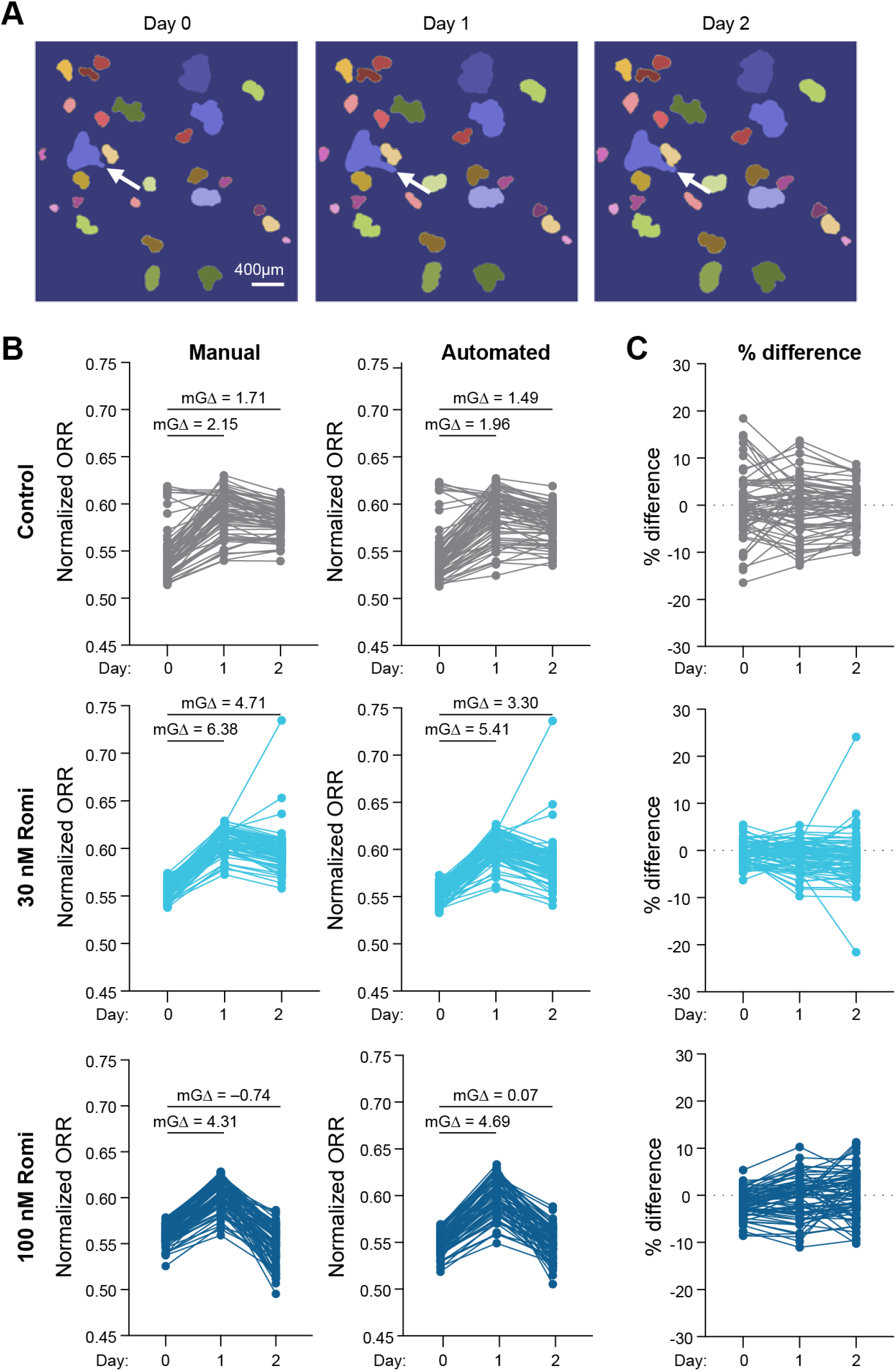
The automated pipeline shows strong agreement with the manual pipeline in tracking single PDCO wide-field redox changes. (A) Representative Nikon images of PDCO-matched masks generated from the tracking algorithm. PDCOs matched across days are highlighted in the same color. White arrows indicate an example of where PDCOs have shifted over time. (B) Keyence ORRs (Eq. 3) are plotted for single PDCOs segmented, tracked, and calculated manually or automatically. mGΔ is used to measure effect size of treatment. (C) Percent differences between manually and automatically calculated ORR are plotted. PDCO numbers for control *n* = 69; 30 nM romidepsin *n* = 79; 100 nM romidepsin *n* = 70.

### Combination of automated segmentation, background normalization, and PDCO tracking accounts for PDCO variability in response

Next, we compared the performance of the entire automated pipeline against our entire manual pipeline in analyzing ORR changes **(Fig. 8)**. The manual pipeline consists of manual segmentation of PDCOs and sampled background normalization of the ORR with no single-PDCO tracking. The novel automated pipeline includes automated segmentation using fine-tuned Cellpose masks, automated background normalization ORR from NAD(P)H and FAD intensity images, and single-PDCO tracking over the treatment time-course.

**Figure 8.**
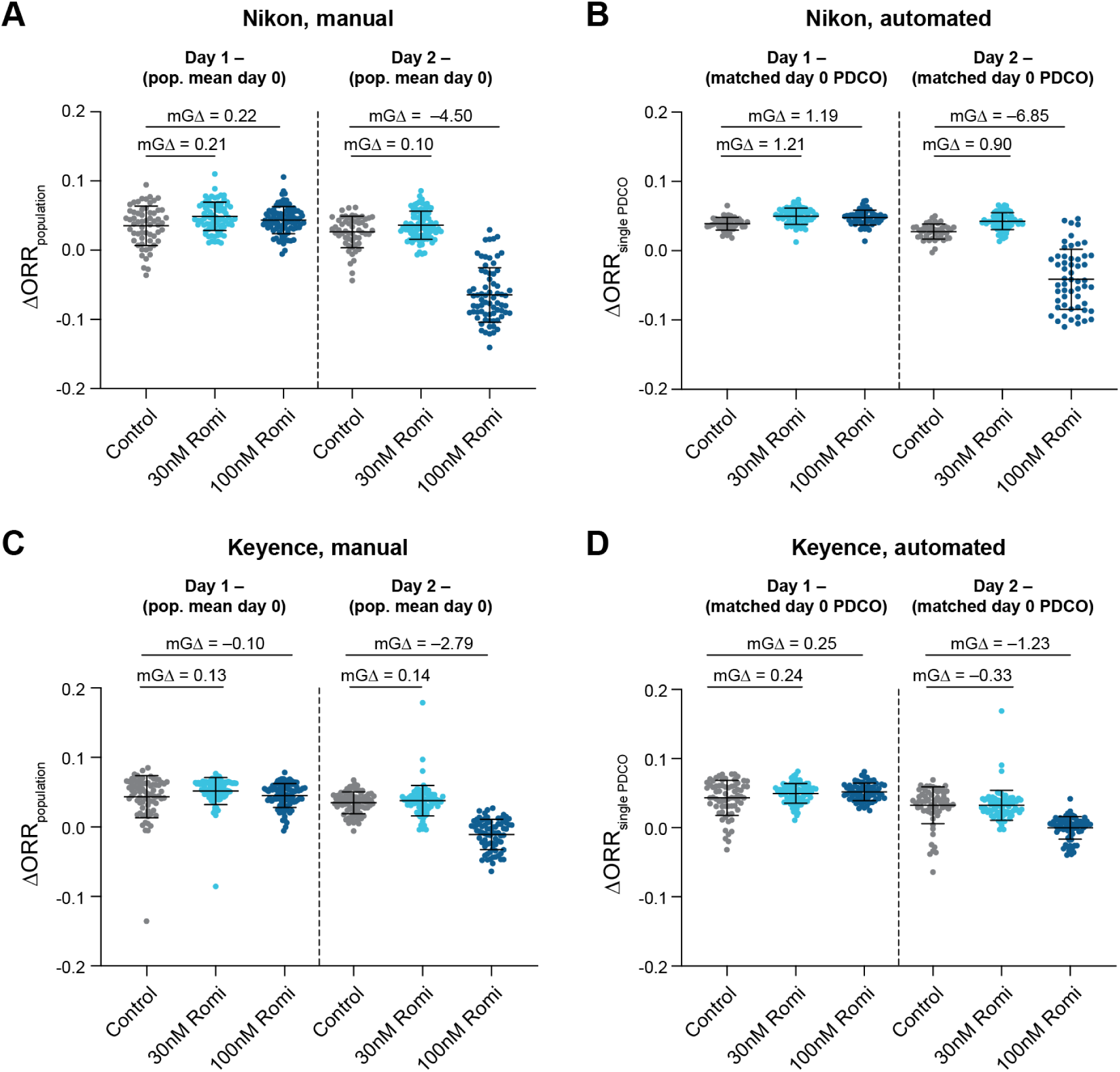
ORR changes obtained with manual and automated pipelines. (A,C) ΔORR_population_ (Eq. 4) were measured using the manual pipeline, which includes manual segmentation and sampled background normalization as previously described, but with no single PDCO tracking, in a (A) Nikon or (C) Keyence microscope. In the manual pipeline, ΔORR_population_ (Eq. 4) was calculated by subtracting the average ORR of a PDCO population before romidepsin (Romi) treatment on day 0 (population day 0) from the ORR of each PDCO in the same population after romidepsin treatment (Romi) on day 1 or 2 as specified. (B,D) ΔORR_single_PDCO_ (Eq. 5) were measured using the entire automated pipeline including segmentation, background normalization, and single PDCO tracking, in a (B) Nikon or (D) Keyence microscope. Here, the ORR of each day 0 PDCO (matched day 0 PDCO) is subtracted from day 1 or day 2 ORR of the same PDCO, excluding unmatched organoids. Number of PDCOs used in (A) and (C) are described in Supplemental Table 1. Supplemental Table 2 describes the number of PDCOs included in single-PDCO tracking analysis (B, D). mGΔ (Eq. 10) is used to measure effect size of treatment. Center line is mean. Error bars are standard deviation.

Here, redox ratio change (ΔORR) is defined differently for the manual and automated pipelines. For the manual analysis ΔORR is calculated using (Eq. 4, ΔORR_population_) by subtracting the average ORR of a PDCO population on day 0 from the ORR of each individual PDCO on days 1 and 2 of treatment. This is due to a lack of tracking capabilities in the manual pipeline. The manual analysis across the Keyence and Nikon systems are consistent, as the day 2, 100 nM romidepsin condition shows the greatest decrease with both microscope systems (**Fig. 8A, 8C**).

We next assessed the performance of the automated pipeline in measuring ΔORR using (Eq. 5, ΔORR_single_PDCO_) where PDCO day 0 ORR is subtracted from day 1 or 2 ORR for each individual PDCO. This is due to the ability to track individual PDCOs over time with the automated pipeline. Notably, almost all standard deviations decreased with single-PDCO tracking (Eq. 5, ΔORR_single_PDCO_) compared to population-level PDCO changes over time (Eq. 4, ΔORR_population_) (**Supplemental Table 3**). This suggests that single-PDCO tracking more accurately accounts for PDCO variability compared to population-level calculations, thus increasing the statistical power of drug response measurements, consistent with prior findings^26^. Comparing across analysis methods, for images acquired on the Nikon, mGΔ increased in all groups with the automated single-PDCO tracking indicating increased sensitivity of the method **(Fig. 8A, 8B)**. For images acquired on the Keyence, mGΔ increased in all groups with the automated single-PDCO tracking except day 2, 100 nM romidepsin condition **(Fig. 8C, 8D)**. Importantly, both microscopes and analysis methods show that the day 2, 100 nM romidepsin group has the largest decrease in ΔORR. Furthermore, the automated algorithm also removes human bias and increases throughput, thus improving the accessibility of wide-field optical redox imaging for testing PDCO drug response.

#### Automated Pipeline Significantly Reduces Analysis Time Compared to Manual Pipeline

We next compared the time required to complete analysis of 9 NAD(P)H and FAD intensity image pairs with the manual pipeline to that of the automated pipeline **(Table 1)**. These images were acquired with the Nikon microscope only. Due to a different FOV size, Keyence images would likely take ∼4 times longer per image for manual analysis. Manual segmentation time is defined as the time between the opening of an image to the time after all PDCOs are segmented. Manual background time is defined as the time between the opening of an image and the time after 5 PDCO-free regions are selected. Manual tracking time was defined as the time required to match IDs for every PDCO across the three NAD(P)H images within a time series, and record the IDs in an Excel sheet, repeated over 3 time-series. In the case of manual tracking, two experts completed the entire dataset independently to ensure accuracy, which required ∼2 hours per three-image time series x 3 time-series (6 hours) per person. The total manual pipeline time includes manual segmentation, manual background, and manual tracking time.

**Table 1:** The time required to complete analysis of 3 sets of 3 time-series images of both NAD(P)H and FAD (9 images for NAD(P)H and 9 images for FAD) with either the manual image analysis pipeline or automated image analysis pipeline. PDCOs were tracked over 3 time points, and images were collected with the Nikon system.

|  | Segmentation Time | Background Time | Tracking Time | Total Pipeline Time |
| --- | --- | --- | --- | --- |
| **Manual** | 1 hour | 9 min | > 6 hours | > 7 hours |
| **Automated** | 37 s | 2.3 min | 25 s | 3.3 min |

The time required to complete analysis of the same 3 sets of 3 time-series images of both NAD(P)H and FAD (9 images for NAD(P)H and 9 images for FAD) with the automated image analysis pipeline was recorded using the Python DateTime module **(Table 1)**. Automated segmentation time was defined as the execution time of the fine-tuned Cellpose model batch processing script. Automated tracking time is defined as the time required to match IDs for every PDCO across the three images **(Fig. 2)**. Automated background time is defined as the execution time of the script that includes the background identification algorithm **(Fig. 3)**. The total automated pipeline time includes automated segmentation, automated background, and automated tracking time.

## Discussion

Wide-field optical redox imaging is a nondestructive approach to assess patient treatment response by tracking longitudinal metabolic changes in PDCOs at the single-PDCO level. This presents an opportunity to apply wide-field optical redox imaging to predict the most effective therapies and identify new therapeutic candidates. To facilitate the expansion of wide-field optical redox imaging as a functional drug screening method in PDCOs, a more accessible and high-throughput image analysis pipeline is needed to meet the demands of large-scale drug screens.

PDCO image analysis is a challenging computational problem due to the variability in PDCO shape, size, and position across focal planes. Autofluorescence images have low SNR, making analysis even more challenging^45^. Several open-source programs, such as OrganoID, MOrgAna, OrgaExtractor, and deepOrganoid, have been developed to address PDCO segmentation and tracking^28–31^. However, these tools are not designed for autofluorescence images, making them unsuitable for assessing wide-field redox imaging data sets.

In some instances, the automated image analysis pipeline developed here disagrees with the manual image analysis pipeline. When comparing between the two pipelines, mGΔ values are similar but not the same. Automated background identification covers a larger area than sampled background identification and likely provides a more accurate representation of the background than only 5 PDCO-free regions selected manually.

Our automated pipeline not only achieves similar sensitivity to drug response in PDCOs as our manual pipeline but also allows us to resolve single-PDCO treatment responses at a significantly faster processing time (>127 times faster, **Table 1**), saving time and limiting user errors. This automated pipeline requires no computational expertise, thereby providing an accessible PDCO image analysis tool. Furthermore, automated single-PDCO tracking enables paired statistical tests that are more sensitive than unpaired population level tests. However, the automated pipeline struggles in some cases. First, because our pipeline relies on dynamic thresholds that account for both position and morphology of PDCOs to track over time, it is less reliable when a PDCO moves or changes in morphology beyond pre-determined thresholds. Second, although the algorithms were designed specifically for autofluorescence images, they still struggle where signals are too low. To improve the generalizability and accuracy of the pipeline, existing deep learning methods may help address these limitations in image variation^28–31^. Third, the automated pipeline is generalizable across the two imaging systems tested (Nikon and Keyence), but image acquisition parameters such as the objective lens used may affect efficacy. Finally, the middle day image (day N), compared to the control image (day N-1), was chosen as the reference for PDCO tracking to minimize the gap between neighboring days on both sides, which increases the probability of matching across all three days. However, if PDCOs are missing due to treatment effects, this choice of reference day could generate errors and therefore the imaging frequency should be increased in those cases.

Overall, this study presents an improved image analysis pipeline that automates PDCO segmentation, single-PDCO tracking, and background correction for wide-field redox imaging. This improves the throughput and robustness of PDCO image analysis and represents a step towards our goal to enhance the accessibility of wide-field optical redox imaging for PDCO drug screening.

## Author Contributions

AH: design, analysis, and interpretation of data, creation of new software used in the work, and drafted the work; KS: design, analysis, and interpretation of data, creation of new software used in the work, and substantively revised the work; AAG: conception, design, analysis, and interpretation of data; SU: acquisition of data; AES: acquisition of data, WZ: analysis of data and creation of new software used in the work; DD: conception and design of the work; MCS: conception, design, and interpretation of data, and substantively revised the work. All authors have read and approved this manuscript.

## Acknowledgements

We thank Skala and Deming lab members for their helpful feedback on the figures and manuscript. We thank Matthew Stefely for his contributions to figure editing. We also thank Dr. Lang Wang for his contributions to the early development of our background algorithm.

## Data/Code availability

Dataset used in this study is available upon reasonable request. The code presented in this study can be found here: https://github.com/skalalab/Auto_WF_PDCO_Pipeline

## Additional Information (Disclosures and Funding Sources)

MCS is an advisor for Elephas Biosciences. This entity had no input in the study design, analysis, manuscript preparation or decision to submit for publication.

The author(s) declare that financial support was received for the research and/or publication of this article. This research was supported by the National Institutes of Health grants R01 CA278051, R01 CA272855, R37 CA226526, Morgridge Institute for Research, Carol Skornicka Chair of Biomedical Imaging. Additionally, the Deming laboratory is funded by the ACI/Schwenn Family Professorship, JD Fluno Family Colorectal Cancer Precision Medicine Program, and the V Foundation.

