## Supplemental Tables for "An automated image analysis pipeline for wide-field optical redox imaging of patient-derived cancer organoids"

**Supplemental Table 1:** Number of PDCOs for each group shown in Fig. 8A, 8C imaged with the Nikon or Keyence system using the manual analysis pipeline.

| | Control (day 0, 1, 2) | 30 nM romidepsin (day 0, 1, 2) | 100 nM romidepsin (day 0, 1, 2) |
|---|---|---|---|
| **Nikon manual** | 59, 60, 56 | 80, 82, 89 | 87, 86, 70 |
| **Keyence manual** | 82, 73, 82 | 91, 88, 90 | 98, 90, 74 |

**Supplemental Table 2:** Number of PDCOs for each group shown in Figures 8B, 8D imaged with the Nikon or Keyence system using the automated analysis pipeline.

| | Control | 30 nM romidepsin | 100 nM romidepsin |
|---|---|---|---|
| **Nikon automated** | 47 | 64 | 55 |
| **Keyence automated** | 69 | 79 | 70 |

**Supplemental Table 3:** Standard deviations for each group shown in Figures 8A-D imaged with the Nikon or Keyence system using either the manual or automated analysis pipelines.

| | **control day 1 - matched day 0 PDCO** | **30 nM Romi day 1 - matched day 0 PDCO** | **100 nM Romi day 1 - matched day 0 PDCO** | **control day 2- matched day 0 PDCO** | **30 nM Romi day 2 - matched day 0 PDCO** | **100 nM Romi day 2 - matched day 0 PDCO** |
|---|---|---|---|---|---|---|
| **Nikon automated** | 0.009 | 0.012 | 0.010 | 0.011 | 0.012 | 0.043 |
| **Keyence automated** | 0.025 | 0.014 | 0.013 | 0.026 | 0.0215 | 0.016 |
| | **control day 1 - population mean day 0** | **30 nM Romi day 1 - population mean day 0** | **100 nM Romi day 1 - population mean day 0** | **control day 2 - population mean day 0** | **30 nM Romi day 2 - population mean day 0** | **100 nM Romi day 2 - population mean day 0** |
| **Nikon manual** | 0.028 | 0.020 | 0.020 | 0.022 | 0.020 | 0.039 |
| **Keyence manual** | 0.030 | 0.019 | 0.017 | 0.015 | 0.0219 | 0.022 |
